# Double-observer approach with camera traps: Towards an unbiased density estimation of unmarked animal populations

**DOI:** 10.1101/2021.04.09.439250

**Authors:** Yoshihiro Nakashima, Shun Hongo, Kaori Mizuno, Gota Yajima, Zeun’s C.B. Dzefck

**Affiliations:** College of Bioresource Science, Nihon University, 1866 Kameino, Fujisawa, Kanagawa 252-0880, Japan; The Center for African Area Studies, Kyoto University, 606-8501, Japan; Projet Coméca, Cameroon

## Abstract

Camera traps are a powerful tool for wildlife surveys. However, camera traps may not always detect animals passing in front. This constraint may create a substantial bias in estimating critical parameters such as the density of unmarked populations. We proposed the ‘double-observer approach’ with camera traps to counter the constraint, which involves setting up a paired camera trap at a station and correcting imperfect detection with a hierarchal capture-recapture model for stratified populations. We performed simulations to evaluate this approach’s reliability and determine how to obtain desirable data for this approach. We then applied it to 12 mammals in Japan and Cameroon. The results showed that the approach could correct imperfect detection as long as paired camera traps detect animals nearly independently (Correlation coefficient < 0.2). Camera traps should be installed to monitor a predefined small focal area from different directions to satisfy this requirement. The field surveys showed that camera trap might miss animals by 3 %–40%, suggesting that current density estimation models relying on perfect detection may underestimate animal density by the same order of magnitude. We hope that our approach will be incorporated into existing density estimation models to improve their accuracy.

## Introduction

Camera trapping is an innovative research technique and has numerous advantages over classic approaches based on human observations. For example, this technique is minimally invasive for animals and habitats and can efficiently collect quantitative records of ground-dwelling mammals, even elusive and nocturnal species, with relatively low labour costs^1–3^. Moreover, by combining it with well-established statistical approaches^4,5^, wildlife occupancy can be reliably estimated^6,7^, abundance or density determined^8–10^, and species richness assessed^11–13^. Furthermore, commercially available camera traps are becoming cheaper, and their battery duration has also been improving^14^, allowing for large-scale surveys in space and time^15^. Data processing (e.g. species identification) may require a long time and heavy labour costs, but a novel technology using machine learning may drastically reduce the cost^16–18^. Given that camera trapping will be increasingly used to make critical decisions on wildlife conservation management, researchers must refine this technique further to avoid arriving at an erroneous conclusion.

One of the primary concerns regarding the application of camera traps may be that they may not always detect animals passing within the camera detection zone (i.e. imperfect detection)^19^. Studies have shown that camera sensitivity may be primarily a function of animal body mass^19,20^ and may also be affected by many factors, such as other animal traits (e.g. typical behaviours), environmental conditions (e.g. temperature and vegetation), and camera efficiency (e.g. trigger speed)^20–27^. This constraint is one of the reasons why the number of animal passes captured by camera traps per unit time (i.e. trapping rate) does not always correlate with animal density and cannot be a reliable index of animal abundance^28^. A series of statistical models are available to account for imperfect detections and reliably estimate animal occupancy and abundance^4^. Nonetheless, researchers cannot help assuming perfect or equal detectability in other situations. In particular, researchers have proposed several analytical approaches to estimate animal density without recognizing animal individuals^29–37^. However, most of these models, except for Chandler and Royle^38^ and Ramsey *et al.*^36^, assume that camera traps can detect animals passing in a specific area within the camera detection zone with absolute certainty (i.e. perfect detection). Rowcliffe *et al.*^19^ suggested that the effective area of the camera detection zone (codeterminants of the camera sensitivity) can be determined using a distance sampling approach by applying detection function models to data on the position where animals were first detected. Although this approach is an important step in quantifying camera sensitivity, it also requires perfect detection at a given point within the camera detection zone, which is still questionable.

A possible yet untested approach to correcting imperfect detection would be to apply the ‘independent double observer approach.’ This approach has been used in point-count surveys based on direct observations by human observers^39–43^. In this approach, two observers (camera traps in our context) record animals concurrently and independently, and the detection probability is estimated from the match or mismatch of observation records. Two analytical models, the capture-recapture and *N*-mixture models can be applied to correct imperfect detections^41^. The former requires that the observers confer with each other regarding each observation, while the latter may be based solely on the counts by each observer. Nichols *et al.*^41^ applied these two models to bird surveys and reported that both models have potential, while the precision of estimates was higher in the former approach. Given that timestamps within images captured by camera traps would allow for efficient reconciling of each count, the capture-recapture model may provide a reliable and efficient means to obtain unbiased estimates of trapping rates.

A constraint in applying the capture-recapture model to the double-observer approach may be the assumption that two observers (i.e. camera traps) detect animals independently. If this assumption is violated, detection probability is overestimated, underestimating the number of animal passes. Regrettably, in practice, the detection probability by paired camera taps may depend on numerous unknown factors (e.g. animal body mass, ambient temperature, etc.), resulting in correlated detections. One viable method to deal with this issue would assume a beta distribution instead of a Bernoulli distribution for the detection processes^44,45^. However, this method may not be a panacea, and thus it is necessary to determine to what extent of the heterogeneity this modelling approach allows for^46^.

This study tested the potential of a hierarchical capture-recapture model using the Bayesian framework to correct imperfect detections and estimate the number of animal passes. Firstly, we introduced the hierarchical capture-recapture model for stratified populations. Secondly, Monte Carlo simulations were performed to evaluate the reliability of this approach, focusing on to what extent the model can accommodate the correlated detections. Thirdly, we performed additional simulations to determine desirable camera selections and placements for this approach. Fourthly, we applied the models to the datasets obtained from two different habitats, Cameroon and Japan, and quantified the detection probability of 12 mammals with varying body sizes.

## Results

### Testing the effectiveness of the hierarchical capture-recapture model

The results of the Monte Carlo simulations showed that the hierarchal capture-recapture models could provide good estimates with a reasonable confidence interval coverage, as long as a pair of camera traps detects a passing animal nearly independently (Table 1). On the other hand, when the correlation coefficient was > 0.2, detection probability was overestimated, underestimating the number of animal passes. This pattern did not differ largely between the lower (0.8) and higher (0.4) detection probabilities (Table 1).

**Table 1.**
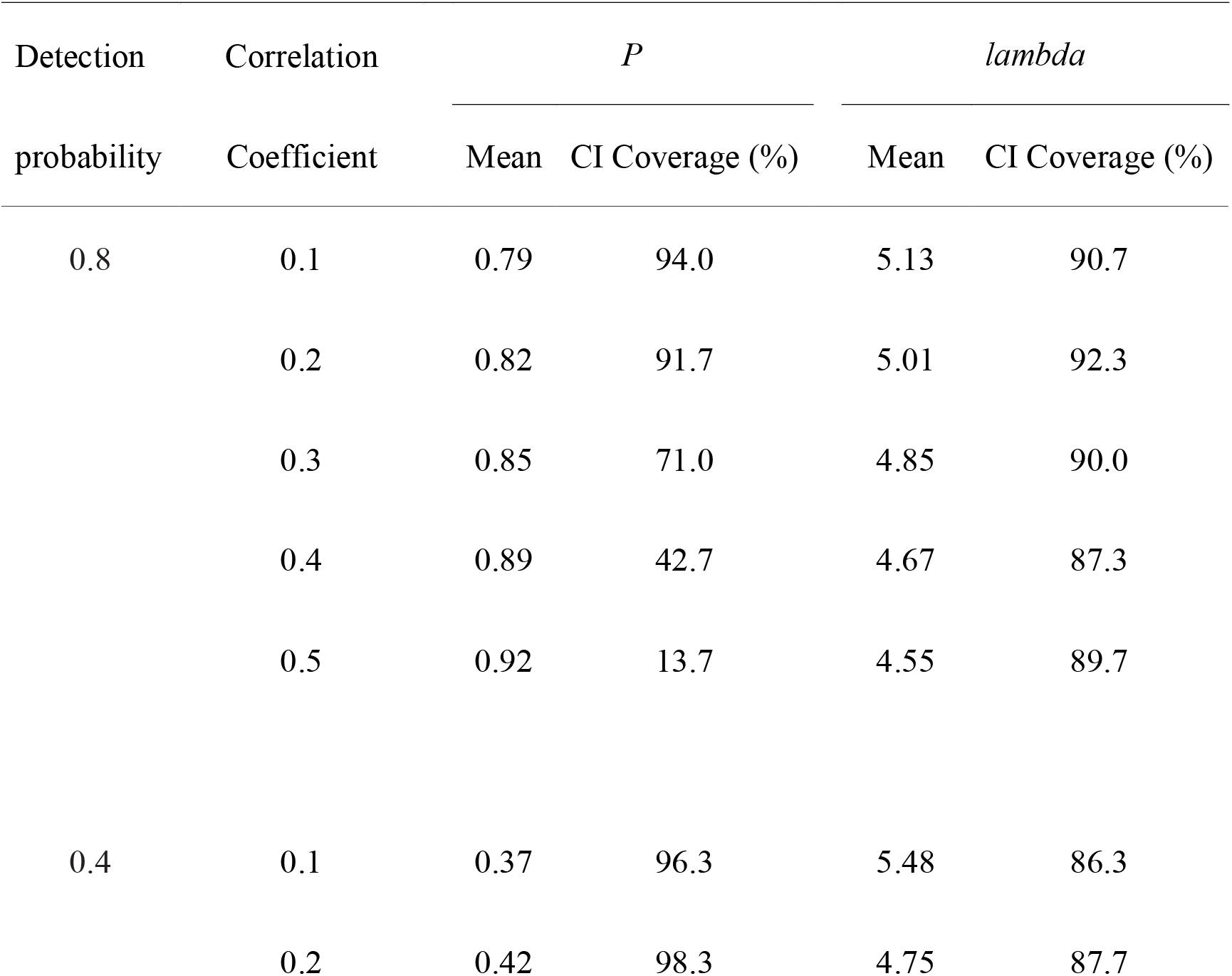
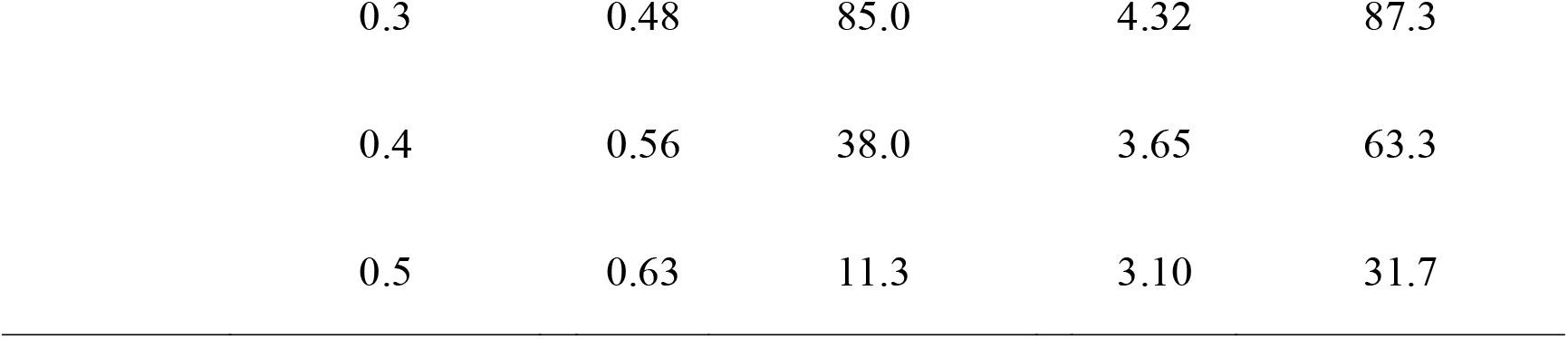
Summary of the Monte Carlo simulations to test the reliability of the hierarchical capture-recapture model. Results are the mean of estimated median detection probability *P,* the expected number of animal passes (lambda), and their 95% credible interval (CI) coverage of the densities. Parameter estimation of the capture-recapture model was performed using the Markov chain Monte Carlo (MCMC) method, and their variances and credible limits were calculated as the posterior summary.

### Determining suitable camera traps and their installations

The simulations mimicking the detection processes of moving animals showed that camera models with a lower trigger speed produced a more correlated detection history (Table 2, Fig. 1). The degree of the correlations also depended on the camera placements. Even when using a camera model with high trigger speed, the detection histories of paired camera traps were highly correlated when monitoring the entire field of view (*r* = 0.53) or the small focal area from the same direction (*r* = 0.27). However, the correlation significantly decreased when monitoring the small focal area from the different directions (*r* = 0.18).

**Table 2.**
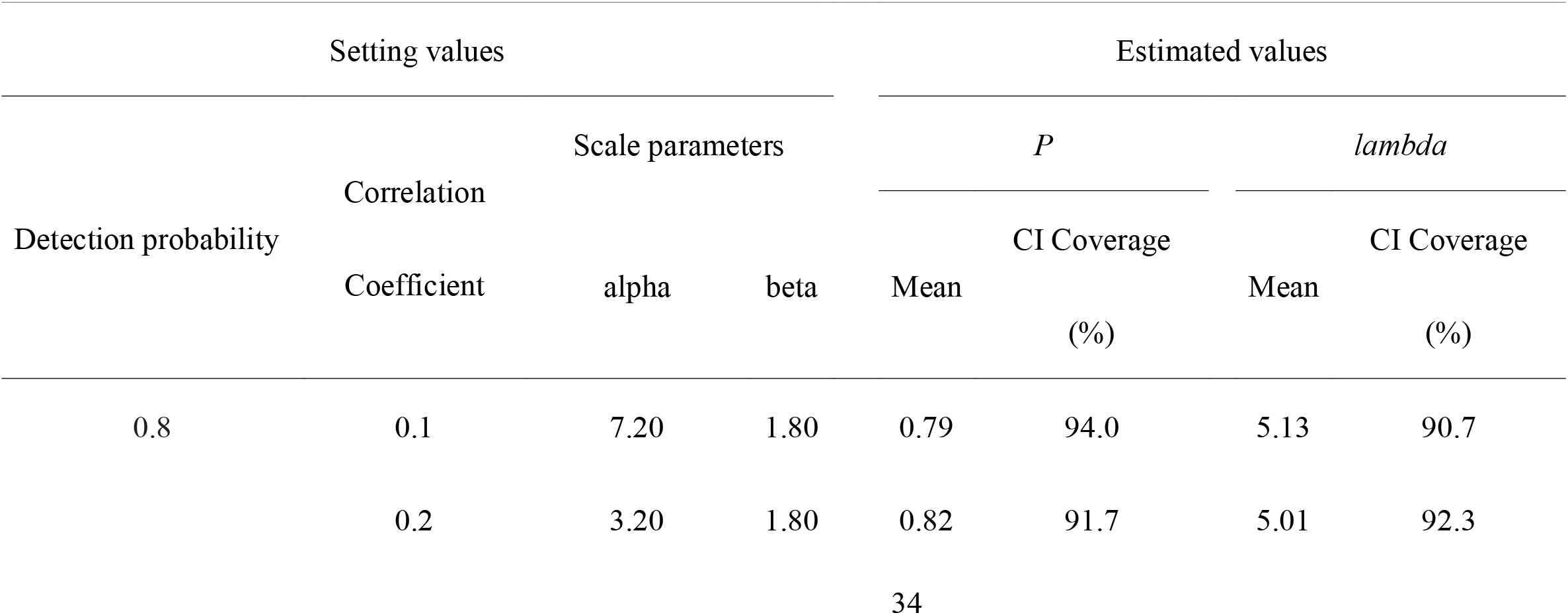
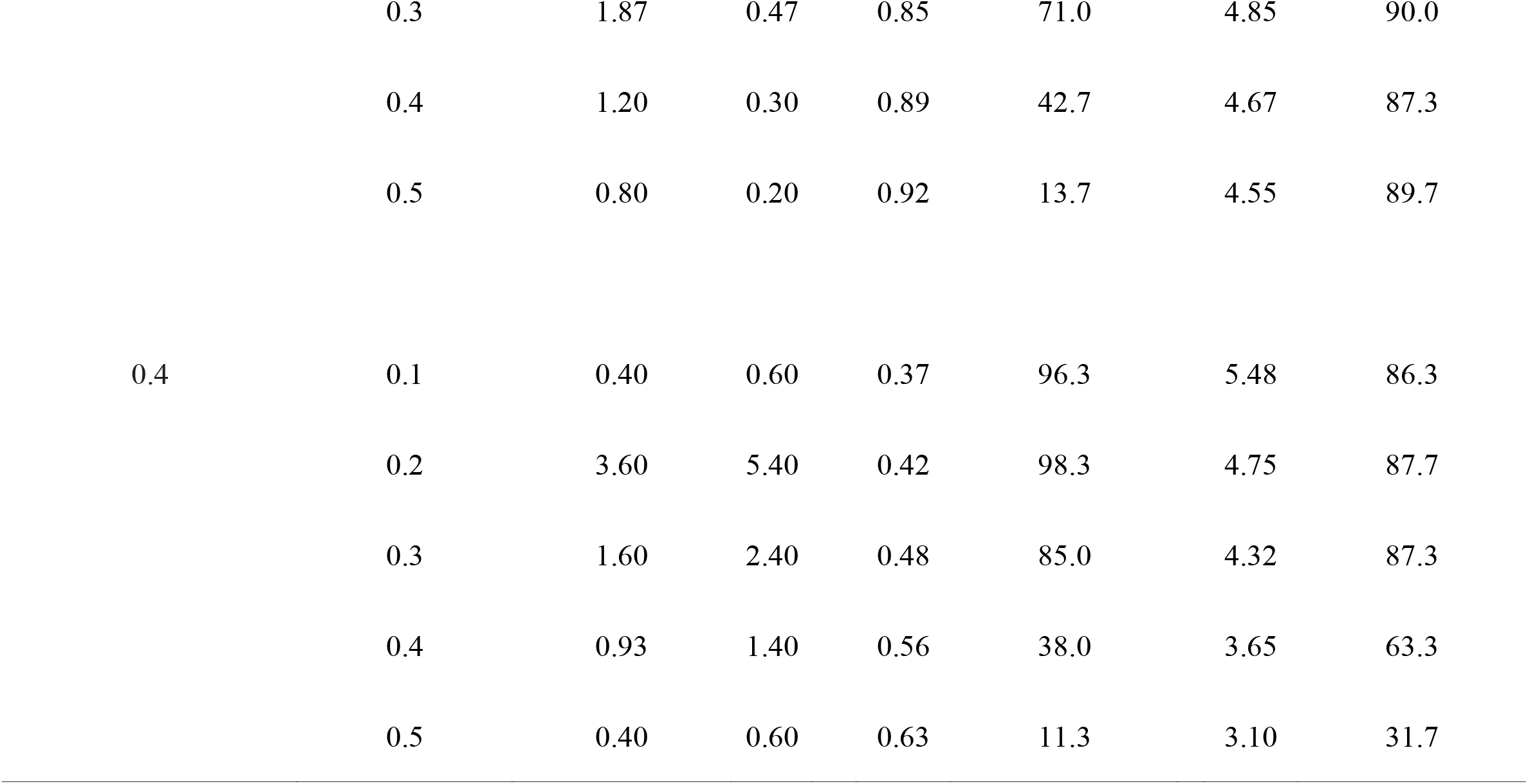
Summary of the Monte Carlo simulations to test the reliability of the hierarchical capture-recapture model. esults are the mean of estimated median detection probability *P* and the expected number of animal passes (*lambda*) and heir 95% credible interval (CI) coverage of the densities. Parameter estimation of the capture-recapture model was erformed using the Markov chain Monte Carlo (MCMC) method, and their variances and credible limits were calculated s the posterior summary.

**Fig. 1.**
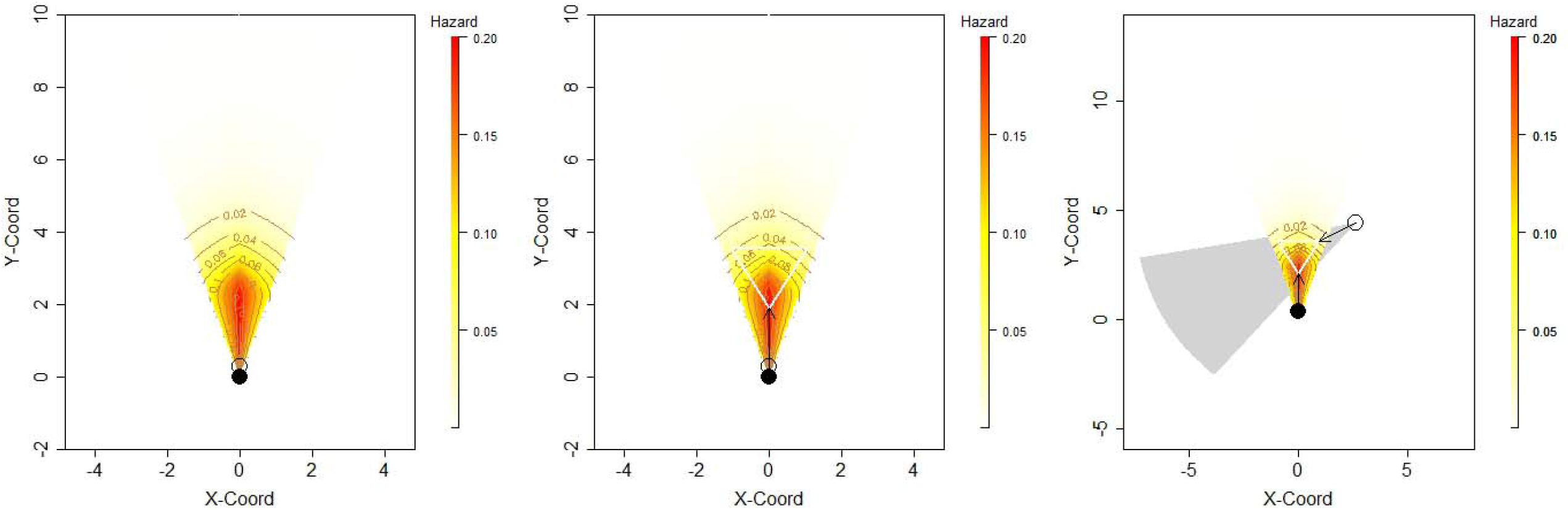
A schematic diagram showing the installations of three focal areas assumed in the simulation mimicking detection processes of moving animals. The circular sector shows the hazard landscape of the detection zone. The open and filled circle shows the positions of camera traps. In ins. 1 (left panel), the focal area was defined as the entire field of view within 10 m from the cameras. In ins. 2 (centre panel), the focal was restricted to an equilateral triangle with a side of 1.9 m. The distance from the camera to the nearest vertex was assumed to be 1.9 m (shown in white lines). Finally, in ins. 3 (right panel), the same equilateral triangle was monitored from different angles of 60 degrees. The camera traps were assumed to have a sensor detection range of 42°. In the right panel, the landscape of the second camera trap was shown in a grey polygon.

### Field surveys

In total, 12 species were recorded more than 10 times (7 in Japan and 5 in Cameroon), which were targets for our analyses. Although none of the species was perfectly detected, the detection probability was estimated relatively high (> 0.8) except for Japanese field mice (95% CI: 0.56-0.64) and tree pangolins (0.62-0.91). The analysis results are summarised in Fig. 2 and Supplementary Table S1.

**Fig. 2.**
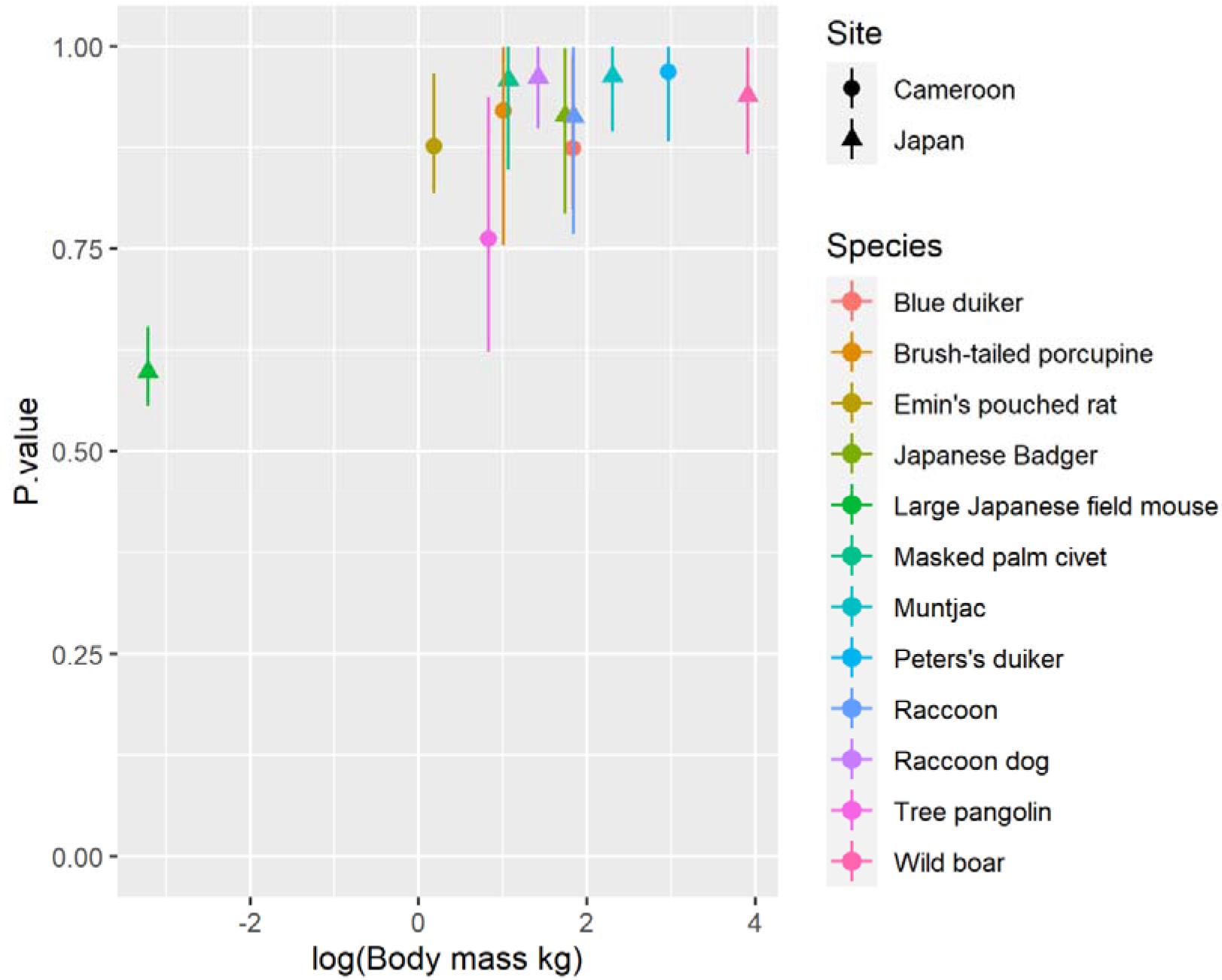
The estimated detection probability of 12 species within a small focal area (1.56 m^2^) in Cameroon (5 species) and Japan (7 species) using the capture-recapture models for stratified populations. Error bar shows the 95% credible interval.

## Discussion

The Monte-Carlo simulation showed that the hierarchical capture-recapture model assuming correlated detections (i.e. using a beta-binomial distribution) provided unbiased estimates of detection probability and the number of animal passes, as long as the correlation coefficients were ≤ 0.2. However, as the correlation increased, the model produced more biased estimates. Regrettably, data obtained from two camera traps may not be sufficient to estimate the parameters of a beta distribution reliably. Indeed, when we assumed installing ten camera traps at each station, the model could provide unbiased estimates even with data having a correlation coefficient of 0.8 (results not shown). Nonetheless, this installation is not practical, and it is more realistic to carefully design a survey to obtain as much independent data as possible.

The simulation mimicking the detection processes of moving animals suggested that camera traps should monitor a small focal area from different directions to obtain nearly independent data. A pair of camera traps facing the same direction have the same hazard landscape, so they often fail to detect animals passing through the periphery of the focal area. On the other hand, camera traps installed in different directions can compensate for weak areas with a lower detection probability by one another. In addition, it may be critical to use a camera model with high trigger speed to avoid missing fast-moving animals. There may be other viable approaches to address heterogeneity in detection probability. For example, one could measure the distance and angle of animals from the camera trap and incorporate them into the model as covariates. This approach is called MRDS (Mark-Recapture Distance Sampling), a well-developed theoretical framework. However, this approach does not always function well^46^ and also requires an intensive field survey. The installations proposed here may be much simpler and could reduce the labour costs.

The results of our field survey showed that it is critical to account for possible imperfect detection in actual analyses. Although the estimated detection probability was relatively high (>0.8) for most species, camera traps could not detect any species completely. In particular, the detection probability was lower for field mice and tree pangolins (Fig. 1), possibly reflecting the smaller body mass (field mice) or the scaly hairs preventing heat radiation from the body interior (tree pangolins). Given that the size and position of the focal area were specified to maximize the detection probability, the sensitivity of the camera model might be less than one throughout the detection zone for these species. This result warns against applying the current density estimation models without accounting for imperfect detection. In the present study, trapping rates were underestimated by 4 %–36%, which led directly to underestimating animal density by the same order of magnitude. Therefore, the double-observer approaches proposed here should be incorporated into the existing density estimation models relying on perfect detections.

We admit that a shortcoming of this approach remains. In particular, it is necessary to have more cameras available, which may be a constraint on implementation. Nonetheless, researchers may be able to challenge these constraints in various ways. For example, installing a pair of cameras at every camera station may not be necessary, as long as the variance in detection probability among camera stations is not too large. Instead, one may choose to install a pair of camera traps in locations with high trapping rates. It may be effective that the study period may be divided into two parts: one in which two cameras are installed to estimate the detection probability and one in which one camera is installed to estimate the trapping rate. Furthermore, if our approach were used at different locations, it would be possible to extrapolate the results to new sites by incorporating environmental conditions and animal characteristics as covariates. Given that many surveys estimating the density of marked populations use paired camera traps at each camera station (to recognize individual animals reliably), it may also be possible to roughly assess the detection probability from currently available data using the approach proposed here.

This study showed that the double-observer approach, combined with hierarchical capture-recapture models using Bayesian frameworks, might be an effective option for estimating detection probabilities and the number of animal passes. We also suggest that commercially available camera traps have higher detection probability within a small focal area but still do not perfectly detect animals. The hierarchical capture-recapture model used here can estimate the distribution of detection probability and the number of animals passing concurrently, and thus, it is readily incorporated into the current density estimation models. We hope that our approach will be incorporated into them to improve their accuracy.

## Methods

### Model framework

The capture-recapture model applied here is the hierarchical model for stratified populations proposed by Royle *et al.*^47^. The model aims to estimate local population size using capture-recapture data from multiple independent locations. In the following, we briefly describe the model in our context, including addressing heterogeneity in detection probability.

Let us consider that we establish *S* independent camera stations in a survey area. Then, we install *K* camera traps at each station to monitor exactly the same focal area (totally *S* × *K* camera traps will be used). We assume that these camera traps detect animals within the focal areas *N*_*T*_ times in total. For animal pass *i* (*i* = 1, 2, 3, ..., *N*_*T*_), we will obtain (1) at which station the animal is detected (hereafter station identity; *g*_*i*_), and (2) how many of the *K* cameras at the station were successful in detecting the animal pass (hereafter detection history; *y*_*i*_). The hierarchal capture-recapture model uses these two vector data, *g*_*i*_ and *y*_*i*_.

Let the number of the animal passes at station *s* be *N*_*s*_ (*s* = 1, 2, 3, ..., *S*). Then, we assume that *N*_*s*_ follows a Poisson distribution with a parameter λ. In this case, the probability of passage *i* occurring at station *s* is expected to be 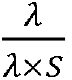. Thus, station identity, *g*_*i*_, can be modelled as follows:

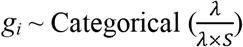

When the number of the animal passes at station *s*, *N*_*s*_, may have larger variation than expected from the Poisson case, we may assume a negative binomial distribution model or may give a random effect to the parameter of the Poisson distribution at the camera station level.

The detection history *y*_*i*_ can be modelled using a data augmentation procedure^47^. Specifically, the original detection *y*_*i*_ is artificially augmented by many *M* – *n* passes with all-zero histories (i.e. not detected by any camera). The augmented data will consist of the passage that occurred but was not detected by any camera (false zero), which occurs with probability *ψ*, and the passage that did not occur (structural zeros) with the probability 1 - *ψ*. A set of latent augmentation binary variables, *z*_*1*_, *z*_*2*_, ... *z*_*M*_, is introduced, which denotes the false zero (*z* = 1) and the structural zero (z = 0). That is

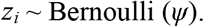

The augmented data *Y*_*i*_ (*y*_*1*_, *y*_*2*_...*y*_*n*_, 0, 0, ... 0) with zero inflation can be modelled conditional on the latent variables *z*_*i*_.

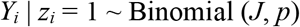

When accounting for the heterogeneity of detection among animal passes, it can be accommodated using a beta distribution as follows;

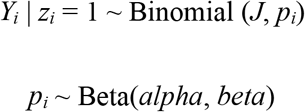

Note that the expected total number of animal passes can be expressed as *λ* × *S*. Thus, *ψ* can be fixed as follows:

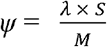

For more details of the models, see Royle *et al.* ^47^.

### Testing the effectiveness of the hierarchical capture-recapture model

We performed Monte Carlo simulations to evaluate the effectiveness of the hierarchical capture-recapture model. Because the model reliability has been confirmed well^48^, we here focused on the effects of heterogeneity in detection probability on the accuracy and precision of the estimates.

We assumed that the number of detections by camera traps followed a negative binomial distribution with a mean of 5.0 and dispersion parameter 1.27, which deprived the actual data on an ungulate in African rainforests^34^. We also assumed two camera traps at 30 stations (i.e. 60 camera traps in total). We generated detection histories (i.e. the number of camera traps successfully detecting animals in each animal passage) using a beta-binomial distribution with the expected detection probability at 0.8 or 0.4. We varied the correlation coefficients (= 1/(alpha + beta + 1),) from 0.1 to 0.5 in 0.1 increments. The scale parameters of the beta distributions for each scenario are shown in Table 1.

We estimated the model parameters using the Markov chain Monte Carlo (MCMC) implemented in JAGS (version 3.4.0) in all the simulations. We assumed that the number of animal passes followed a negative binomial distribution, and the detection probability did a beta distribution. Since the estimation of scale parameters (*α* and *β*) of the beta distribution, we transformed as *α* = *p*\**phi* and *β* = *p*\*(1 - *phi*) (*p* is an expected detection probability). Then we used weakly informative prior (gamma distribution with shape = 10 and rate = 2) for *phi* while a non-informative uniform distribution from 0 to 1 for the detection probability^49^. We generated three chains of 3000 iterations after a burn-in of 1000 and thinned by 5. The convergence of models was determined using the Gelman–Rubin statistic, where values <1.1 indicated convergence. These procedures were repeated 300 times. The JAGS code is available as supplementary material (Supplementary R1).

### Determining a suitable camera installation

The above simulations suggested that a key to safely applying the hierarchical capture-recapture model may avoid correlated detections. To determine a preferred survey design to secure independent detections, we performed additional simulations. Specifically, we tested how the position and the trigger speed of camera traps may affect the independence of detections by considering the process by which cameras detect moving animals.

The simulation was performed using similar procedures taken by Rowcliffe *et al.* ^19^. We assumed that the sensor of camera traps has a two-dimensional detection surface defining the instantaneous ‘risk’ (as analogous to the risk of mortality in survivorship analysis) of an animal being detected at any given location within the camera’s field of view (FOV). The instantaneous risk landscape was defined with respect to distance *r* and angle relative to the camera as follows:

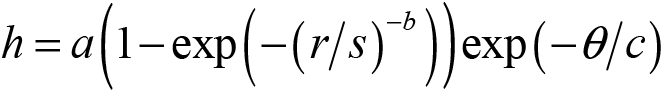

where *a* defines the maximum risk close to the sensor, *s* and *b* define the position and shape of decline in risk with distance, respectively, and *c* defines the rate of decline in risk with angle. We set *a*, *s*, *b*, and *c* at 0.2, 3.0, 5.0, and 0.5, respectively. Although there is no empirical evidence for this particular function form, the simulated results were comparable to those observed. Note that our interest is not in estimating the actual values of the correlation but instead in obtaining information that will help us decide what cameras to install and how to install them.

We made an animal pass through the risk landscape in a straight line in a random direction. The animal movement speed (ms^−1^) followed a log-normal distribution with a mean (± *SD*) of 0.18 ± 0.15 m, which roughly accord with the speed of red duikers in our study sites (Y Nakashima, unpublished data). We then generated a random animal’s position to be detected, considering the cumulative risk of detection and the camera trap’s trigger speed (for the details, see Rowcliffe *et al.*^19^).

We considered three designs of camera placements (Fig. 1). The first one is to install two camera traps at the same position (i.e. mounted on the same tree) and in the same direction to monitor the entire field of view within 10 m from the cameras (ins. 1). The second and third one is assumed to monitor a specific small area within the FOV. The focal area was a small equilateral triangle with a side length of 1.9m and was centred within the FOV. The nearest vertex was set to be 1.9m away from the cameras. This area corresponds to the highest detection probability in the camera model used in our field study. This focal area is monitored from the same direction (ins. 2) or different angles of 60 degrees (ins. 3). We then considered using camera models with a fast trigger speed (0.1 s) and a slow (1.5 s) for each installation. Finally, we generated the detection history of 500 times animal passes and calculated the correlation coefficients between the two camera traps for each scenario.

### Field surveys

We conducted a field survey in and around the Boumba-Bek and Nki National Parks in southeast Cameroon (October 2018 to January 2019) and the Boso Peninsula in Japan (August-November 2018). The study area of Cameroon consists primarily of evergreen and semi-deciduous forests. The annual rainfall is approximately 1,500 mm, and the mean annual temperature is *ca*. 24°C. Typically, the dry season occurs from December to February and the rainy season is from March to November. The Boso Peninsula was in the southern part of Chiba Prefecture in central Japan (35N °N, 140E). The vegetation consists of either broad-leaved evergreen forests (*Castanopsis sieboldii* and *Quercus* spp.) or coniferous plantations (*Cryptomeria japonica* and *Chamaecyparis obtusa*). The monthly mean temperature was 20.6 ± 5.9 °C during the study period, with the highest in August (27.4 °C) and the lowest (13.6 °C) in November.

The simulations suggest that camera traps monitor the predefined focal area from different positions (see below). According to the results, we used the camera traps with a high trigger speed (0.15 s) (Browning Strike Force Pro, BTC-5HDP, Browning, Missouri, US) at both study sites. We regarded two camera traps to monitor the same equilateral triangle area as in the simulation from different directions by 60 degrees (the right panel of Fig. 1). We surrounded the focal area with a white rope and manually filmed it with camera traps as a reference. The rope was removed after filming to avoid disturbing the animal behaviour. We set camera traps at approximately 0.7 m above ground without baits or lures. We used the ‘video mode’ and designated the video length as 20 s and the delay period between videos at 1 s (minimum delay period in this product). In Japan, we established seven camera stations at least 2 km apart and installed two camera traps at each station. In Cameroon, we set 26 camera stations at least 2-km apart from each other. Since the number of camera stations is not enough to estimate the expected number of animal passes in Japan, we focused on the detection probability

We determined whether animals passed within or outside the focal area by superimposing videos and the reference image. We used only images of animals crossing the focal area for subsequent analyses. We then matched each detection from the two cameras to determine whether two camera traps successfully recorded an animal pass. We applied the model to the detection probability of the species detected more than 10 times in both Japan and Cameroon. The analysis was limited to images in which the animal species were reliably identified. We plotted the estimated detection probability against the median value of body mass (kg), drawn from Ohdachi et al.^50^ for animals in Japan and Kingdon^51^ for those in Cameroon.

## Acknowledgements

This study complied with the Republic of Cameroon laws. It was conducted with approval from the Ministry of Scientific Research and Innovation (MINRESI, N°0190/MINRESI/Projet COMECA/PM/07/2018) and the Ministry of Forestry and Wildlife (MINFOF, N°1527/L/MINFOF/SETAT/SG/DFAP/SDCF/SEP/EP), Cameroon. This study was financially supported by JST/JICA SATREPS (JPMJSA1702) and JSPS KAKENHI Grant Number 15K07487 and 21H03653.

## Contributions

YN conceived the ideas and designed the methodology; SH and YG collected the data; KM and YG performed video analyses; YN wrote the manuscript. All authors contributed critically to the drafts and approved the final manuscript for publication.

## Ethics declarations

### Data availability

All the data used here is available in Supplementary Information.

