## Supplementary Table S1 and Supplementary R code for "Double-observer approach with camera traps: Towards an unbiased density estimation of unmarked animal populations"

| Supplementary Table S1. Summary of the applications of the hierarchical capture-recapture model to the field data in Japan and Cameroon. Results are the number of the animal passes detected (*N*); Pass: animal's pass specific random effects), the mean and median of estimated detection probability *P* and the 95% credible interval. Parameter estimation of the capture-recapture model was performed using the Markov chain Monte Carlo (MCMC) method, and their variances and credible limits were calculated as the posterior summary. Body mass (kg) was adapted from Kingdon *et al.* (2015) or Ohdachi *et al.* (2009). | | | | | | | | |
| --- | --- | --- | --- | --- | --- | --- | --- | --- |
|  |  |  | Body mass |  | *P* | | | |
| Site | Species | Scientific name | (kg) | *N* | Median | 95% CI | | |
| Cameroon | Blue duiker | *Philantomba monticola* | 6.3 | 78 | 0.87 | 0.80 | - | 0.97 |
|  | African brush-tailed porcupine | *Atherurus africanus* | 2.8 | 13 | 0.92 | 0.75 | - | 1.00 |
|  | Emin's pouched rat | *Cricetomys emini* | 1.2 | 140 | 0.88 | 0.82 | - | 0.95 |
|  | Peters's duiker | *Cephalophus callipygus* | 19.5 | 54 | 0.97 | 0.88 | - | 1.00 |
|  | Tree pangolin | *Phataginus tricuspis* | 2.3 | 24 | 0.76 | 0.62 | - | 0.91 |
| Japan | Wild boar | *Sus scrofa* | 50 | 117 | 0.94 | 0.87 | - | 0.99 |
|  | Japanese Badger | *Meles anakuma* | 5.7 | 33 | 0.91 | 0.79 | - | 0.99 |
|  | Large Japanese field mouse | *Cricetomys emini* | 0 | 279 | 0.60 | 0.56 | - | 0.64 |
|  | Masked palm civet | *Paguma larvata* | 2.9 | 27 | 0.96 | 0.85 | - | 1.00 |
|  | Muntjac | *Muntiacus reevesi* | 10 | 67 | 0.96 | 0.89 | - | 1.00 |
|  | Raccoon | *Procyon lotor* | 6.3 | 23 | 0.91 | 0.77 | - | 1.00 |
|  | Raccoon dog | *Nyctereutes procyonoides* | 4.1 | 127 | 0.96 | 0.9 | - | 1.00 |

Supplementary R1. R code to perform the Monte-Carlo simulations

\# to test the reliability of hierarchical capture-recapture models for a stratified population

\# to estimate detection probability and correct potentially imperfect detection.

library(MASS)

library(jagsUI)

library(extraDistr)

nsim <- 3 # Number of iteration

nstation <- 30 # Number of camera stations

ncamera <- 2 # Number of cameras at a station

lambda <- 5 # Parameter for trapping rate

theta <- 1.27 #1.27 #1.27 # Dispersion parameter for trapping rate

expp <- 0.8 # Parameter for detection probability

cor <- 0.3 # Correlation coefficient

neg_beta_p <- neg_beta_lambda <- list(0)

t <- proc.time()

simn <- 0

for (m in 1:nsim) {

simn <- simn + 1

cat(paste(simn, ", ", sep = ""))

npass_station <- rnegbin(nstation, lambda, theta = theta)

npass_total <- sum(npass_station)

station_pass <- rep(1:nstation, npass_station)

\# Beta model without random effects

pbeta <- expp

phi <- 1/cor - 1

(alpha <- pbeta * phi)

(beta <- (1 - pbeta) * phi)

\# hist(rbeta(10000, alpha, beta))

y.full <- rbbinom(npass_total, ncamera, alpha = alpha, beta = beta)

y <- y.full[y.full != 0]

w <- c(y, rep(0, length(y) * 1.2)) # Augmented detection history

M <- length(w)

Q <- length(y)

V <- station_pass[y.full != 0]

g <- c(V, rep(NA, M - Q))

K <- ncamera

datalist <- list(w = w, nstation = nstation, M = M, K = K, g = g)

\# Beta model --------

model <- "C:\\bugstemp\\model_HCR_NB.txt"

sink(model)

cat("

model {

p ~ dunif(0,1)

alpha.lam ~ dgamma(0.01, 0.01)

psi <- sum(lam[])/M

eta ~ dunif(0,100)

phi ~ dgamma(10,2)

for(s in 1:nstation){

log(lam[s])<-log(alpha.lam) + log(eps[s])

gprobs[s]<-lam[s]/sum(lam[])

eps[s] ~ dgamma(eta, eta)

}

for(i in 1:M){

g[i] ~ dcat(gprobs[])

z[i] ~ dbern(psi)

w[i] ~ dbin(mu[i], K)

mu[i] <- z[i]*alpha.p[i]

alpha.p[i] ~ dbeta(p*phi,(1-p)*phi)T(0.01,0.99)

}

theta <- 1- 1/phi

}

",

fill = TRUE)

sink()

inits <- function() list(z = rep(1, M))

parameters <- c("alpha", "beta", "theta", "p", "psi", "alpha.lam")

n.chain <- 3

n.iter <- 3000

n.burnin <- 1000

n.thin <- 5

nmcmc <- n.chain * (n.iter - n.burnin)/n.thin

neg_beta <- jags(datalist, inits, parameters, model, n.chain = n.chain, n.iter = n.iter, n.burnin = n.burnin, n.thin = n.thin, parallel = TRUE,

modules = "mix")

neg_beta_p[[m]] <- c(neg_beta$q50$p, neg_beta$q2.5$p, neg_beta$q97.5$p)

neg_beta_lambda[[m]] <- c(neg_beta$q50$alpha.lam, neg_beta$q2.5$alpha.lam, neg_beta$q97.5$alpha.lam)

}

par(mfrow = c(2, 1))

par(mar = c(3, 4, 3, 2) + 0.1)

par(mgp = c(2.2, 1, 0))

lam.beta <- do.call(rbind, neg_beta_lambda)

plot(1:nsim, lam.beta[, 1], ylim = c(0, 10), main = "lambda beta")

segments(1:nsim, lam.beta[, 2], 1:nsim, lam.beta[, 3], col = 2)

lines(c(-1, nsim * 2), c(lambda, lambda), lty = 2)

p.beta <- do.call(rbind, neg_beta_p)

plot(1:nsim, p.beta[, 1], ylim = c(0, 1), main = "p beta")

segments(1:nsim, p.beta[, 2], 1:nsim, p.beta[, 3], col = 2)

lines(c(-1, nsim * 2), c(expp, expp), lty = 2)

(proc.time() - t)/60
